# A moonlighting serpin of *Tannerella forsythia* compromises the bactericidal activity of neutrophils

**DOI:** 10.64898/2026.10.08.756777

**Authors:** Savannah C Morris, Florian Veillard, Danuta Mizgalska, Mariusz Madej, Malgorzata Benedyk-Machaczka, Justyna Folkert, Barbara Potempa, Andrew Fuchs, Jan Potempa, Juhi Bagaitkar

## Abstract

Serine protease inhibitors (serpins) are extremely rare in prokaryotes, and their role in immune evasion remains unclear. We identified a bifunctional TLR2-binding serpin (miropin) produced by *Tannerella forsythia* that broadly suppresses neutrophil antimicrobial capacity. Miropin binding to TLR2 primed neutrophils, enhancing phagocytic capacity yet, counterintuitively, promoting bacterial persistence. Mechanistically, like other serpins, miropin acted as an irreversible suicide inhibitor of neutrophil serine proteases (NSPs), blocking their activity upon granule mobilization and NETosis. Unexpectedly, miropin, independent of its protease-inhibitory activity, bound and inhibited myeloperoxidase (MPO), thereby blocking the oxidative burst. This simultaneous suppression of oxidative and non-oxidative bactericidal mechanisms promoted bacterial survival in vivo without significantly altering the tissue inflammatory response. Thus, we identified novel moonlighting roles (TLR2 signaling and MPO inhibition) for an unorthodox prokaryotic serpin and provided context for how *T. forsythia* and other oral pathogens survive within the subgingival niche, an environment highly enriched in activated neutrophils.

---

Neutrophils are continuously recruited to the oral mucosa, where they play an unparalleled role in antimicrobial surveillance and in maintaining oral immune homeostasis. They lead the initial response against several oral pathogens, rapidly phagocytosing and degrading bacteria within phagosomes ^1^. In neutrophils, the phagosomal antimicrobial capacity relies heavily on the activation of the NADPH oxidase (NOX2)-mediated oxidative burst and downstream myeloperoxidase (MPO) activity, which consumes hydrogen peroxide (H_2_O_2_) to generate cytotoxic hypohalous acids ^2^. In addition, trafficking of neutrophil granules and other vesicles to the phagosome delivers additional antimicrobial cargo, such as antimicrobial peptides and neutrophil serine proteases (NSPs) that degrade bacterial membrane proteins ^3,4^, thereby killing ingested bacteria within minutes. However, in pathogenic conditions such as periodontitis, despite increased recruitment and activation of neutrophils in the oral cavity, oral dysbiosis persists and, counterintuitively, is accompanied by an enrichment of certain oral pathobionts that resist neutrophil-mediated clearance through mechanisms that remain incompletely understood.

*Tannerella forsythia* is an understudied bacterial pathogen that is often enriched in the dysbiotic oral microbiome of patients with periodontitis ^5–7^. Similar to other periodontal pathogens (*Porphyromonas gingivalis, Treponema denticola*, *and Fusobacterium nucleatum*), *T. forsythia* is asaccharolytic and synthesizes more than 20 proteases, which it uses to degrade host tissues and obtain nutrients from the resulting breakdown products ^8,9^. Furthermore, studies in animal models demonstrate that *T. forsythia* virulence factors (BspA, lipoproteins) elicit strong inflammatory responses and neutrophil recruitment, leading to skin abscesses in rabbits and mice, as well as alveolar bone loss in mice and rats ^6,10,11^. Thus, to persist in these immune-activated environments, *T. forsythia* requires strategies to counter the host’s antimicrobial arsenal and evade immune clearance. However, despite the clinical association of *T. forsythia* with periodontitis and its ability to induce inflammatory bone loss (alveolar bone recession), very little is known about how it evades immune clearance in the periodontal pocket, a highly inflammatory environment dominated by neutrophils. Here, we focused on ‘miropin’, an atypical prokaryotic **ser**ine **p**rotease **in**hibitor (serpin), to counteract neutrophil antimicrobial action.

Serpins constitute a large family of structurally conserved protease inhibitors found ubiquitously in higher multicellular eukaryotes, playing an important role in the homeostatic regulation of the complement and coagulation proteolytic cascades, thereby limiting tissue damage in humans. However, their presence in prokaryotes is sporadic, and their roles in immune evasion and pathogenesis are poorly understood ^12^. We recently identified ‘miropin’, a novel and highly unusual serpin produced by *T. forsythia*, that inhibits a wide range of mammalian proteases such as trypsin, plasmin, and multiple NSPs ^13–15^. Functionally, serpins use an exposed, flexible stretch of amino acids, called the reactive site loop (RCL), as the primary functional domain for protease inhibition. A protease attack on the active-site peptide bond (P1-P1’) triggers a dramatic structural change in which the RCL is incorporated into the serpin’s large beta-sheet. This pulls the covalently attached protease to the opposite side of the serpin, distorting it into an irreversibly inactive conformation ^16^. For miropin, this broad spectrum of targeting of multiple proteases stems from multiple active sites within the reactive center loop (RCL) and an adaptable protein core, enabling effective targeting of diverse proteases ^13,15^. Thus, we hypothesized that miropin would play an important role in immune subversion during oral mucosal inflammation, where activation of multiple host proteases is essential to limit bacterial invasion.

In this mechanistic study, we show that miropin serves as a versatile virulence factor that simultaneously disengages multiple neutrophil antimicrobial pathways and is essential for *T. forsythia* survival in vitro and in vivo. We show that miropin engages TLR2 to prime neutrophils but does not induce full activation. Instead, miropin, consistent with its serpin function, utilizes its RCL domain to inhibit NSPs while its core binds and inhibits MPO with high affinity, attenuating the phagosomal oxidative burst. The immunosuppressive effect of miropin was not limited to *T. forsythia*, as neutrophils exposed to miropin also failed to clear other pathogens. Overall, our studies substantiate the novel role of miropin as an exceptional triple-specificity protein that functions as a typical protease-inhibiting serpin, a TLR2 ligand, and an MPO-inhibitory protein, the first of its kind among known serpins regardless of their phylogenetic origin.

## METHODS

### Mice

Male and female C57BL/6J and B6.Cg-Tyrc-J/J mice (8-12 weeks) were purchased from The Jackson Laboratory and housed under specific-pathogen-free conditions. All animal experiments were approved by the Institutional Animal Care and Use Committee at The Abigail Wexner Research Institute at Nationwide Children’s Hospital.

### Neutrophil isolation

Neutrophils were isolated from de-identified venous blood obtained from healthy donors, as previously described ^17,18^. Briefly, whole heparinized blood was mixed with 3% (w/v) dextran (MW: 450,000-650,000) to sediment RBCs. Leukocyte-rich plasma was collected, subjected to hypotonic RBC lysis, and neutrophils were isolated by density gradient sedimentation using Histopaque-1077 (Sigma). Purity was assessed by cytospins and Wright-Giemsa staining.

### Bacteria

*Tannerella forsythia* ATCC 43037 (WT *Tf*) was cultured in enriched tryptic soy broth (eTSB; Millipore Sigma) supplemented with 25 g/L yeast extract, 5% horse serum (Gibco), 0.25 g/L L-cysteine, 1 mg/L hemin, 0.5 mg/L menadione, and 10 mg/L N-acetylmuramic acid. The isogenic miropin-null mutant (Δ*mir Tf*) ^14^ was cultured under identical conditions with chloramphenicol (10 µg/ml) selection. *Porphyromonas gingivalis* ATCC 33277 (*Pg*) was grown in eTSB supplemented with 5 g/L yeast extract, 5 mg/L hemin, and 2 mg/L menadione. Bacterial cultures were maintained in an anaerobic chamber (Coy Laboratory Products) containing 92% N_2_, 5% CO_2_, and 3% H_2_ at 37°C. Solid media for both bacteria utilized the aforementioned growth media with 1.5% agar and 5% sheep blood for *Pg*. *Staphylococcus aureus* USA300 (*Sa*) was cultured aerobically in tryptic soy broth (TSB) at 37°C. For all experiments, bacteria were harvested during the logarithmic growth phase at optical densities (OD600) of 0.4-0.6 for *Tf*, 0.8-1.0 for *P. gingivalis*, and 1.0-2.0 for *S. aureus*.

### Protease inhibition assays

Proteases were incubated with a 10-fold molar excess of recombinant miropin variants or alpha-1 anti-trypsin (α1-AT; Sigma-Aldrich) in black 96-well plates in a total volume of 180 µl of 0.1 M Tris-HCl, 150 mM NaCl, 5 mM EDTA, and 0.02% Tween-20, pH 7.6 (except for trypsin assays, which contained 5 mM CaCl_2_ in place of NaCl) for 30 minutes at 37°C. Residual protease activity was measured after adding 20 µl of fluorogenic substrates in DMSO (final concentration, 50 µM): Z-FR-AMC for trypsin, AAPF-AMC for cathepsin G, and AAPV-AMC for human neutrophil elastase. Fluorescence was monitored using a SpectraMax Gemini XS plate reader (Molecular Devices) at excitation/emission of 350/460 nm. Final enzyme concentrations were 5 nM for trypsin and 10 nM for cathepsin G and human neutrophil elastase.

### Quantification of phagosomal uptake, acidification, and MPO activity

Neutrophils were first loaded with 5 µM APF for 30 min at 37°C and then suspended in RPMI supplemented with 2% FBS at 2 x 10^6^ cells/ml. Cells were incubated with 2 µM rMir^WT^ or rMir^G4^ for 30 min at 37 °C and then challenged with pHrodo-labelled *P. gingivalis* or *Tf* (10 bacteria; 1 cell) or with 0.1mg/mL pHrodo Zymosan (Thermo Fisher) for 15, 30, or 60 min at 37°C. Reactions were terminated by adding ice-cold PBS and immediately transferring to ice. Cells were transferred to flow buffer and analyzed by flow cytometry or imaging flow cytometry. In certain experiments to block phagocytic receptors, cells were pre-incubated with anti-TLR2 neutralizing antibody (clone mab2-mtlr2; InvivoGen) for 1 h at 37 °C prior to rMir treatment. Relative phagocytic capacity was measured as the % of pHrodo particles at each time point. Mean fluorescence intensity (MFI) of pHrodo dye and APF was recorded as a measure of phagosomal acidification and MPO activity, respectively. To determine receptor abundance on neutrophils after miropin treatment, neutrophils were treated with 1 µM rMir^WT^ for 1 h. Reactions were stopped with ice-cold FACS buffer, and cells were blocked with 10% human AB serum (except when CD16b antibodies were present). Cells were stained with antibodies against CD11b, CD16b (clone CLB-gran11.5; BD Optibuild), CD35, CD62L (clone DREG56; Invitrogen), CD63, TLR2 (CD282) (clone TLR2.1; Biolegend), and TLR4 (CD284) (clone HTA125; Biolegend) for 1 h on ice in the dark.

### Statistical analysis

All data are presented as mean ± standard deviation (SD) unless otherwise stated. Statistical analyses were performed using GraphPad Prism (v5.0d). Comparisons between two groups were performed using an unpaired Student’s *t*-test or Welch’s t-test. Comparisons involving three or more groups were analyzed using one- or two-way ANOVA, as appropriate. Statistical significance was defined as *p* ≤ 0.05. Specific statistical tests and multiple-comparison procedures are indicated in the corresponding figure legends.

All other methods are listed in the Supplemental Information

## RESULTS

### Miropin is essential for Tf survival within neutrophils

To establish miropin’s role in immune evasion, we first determined whether miropin is essential for *T. forsythia* survival within human neutrophils and then confirmed our findings in three independent *in vivo* murine models. While the wild-type *T. forsythia* (*Tf* WT) strain persisted within human neutrophils, the congenic miropin-deficient mutant (*TfΔmir*) ^14^ was rapidly cleared (**Figure 1A**), indicating that miropin was essential for resisting neutrophil-mediated clearance. Next, to determine the relevance of miropin in the oral cavity, a natural niche for *Tf*, we orally colonized mice with *Tf* WT and *TfΔmir*. While *Tf* WT led to modest yet significant alveolar bone recession (**Figure 1B, C**), the *TfΔmir* strain failed to colonize the oral cavities of mice, making direct comparisons between the strains and their ensuing inflammatory response unfeasible. Thus, to directly assess the role of miropin in *Tf* survival, we employed two independent *in vivo* models (bacterial peritonitis^19–21^ and thigh abscess^22^) in which bacterial introduction leads to rapid neutrophil recruitment and accumulation, allowing quantitative assessment of neutrophil-mediated bacterial clearance *in vivo*.

**Figure 1:**
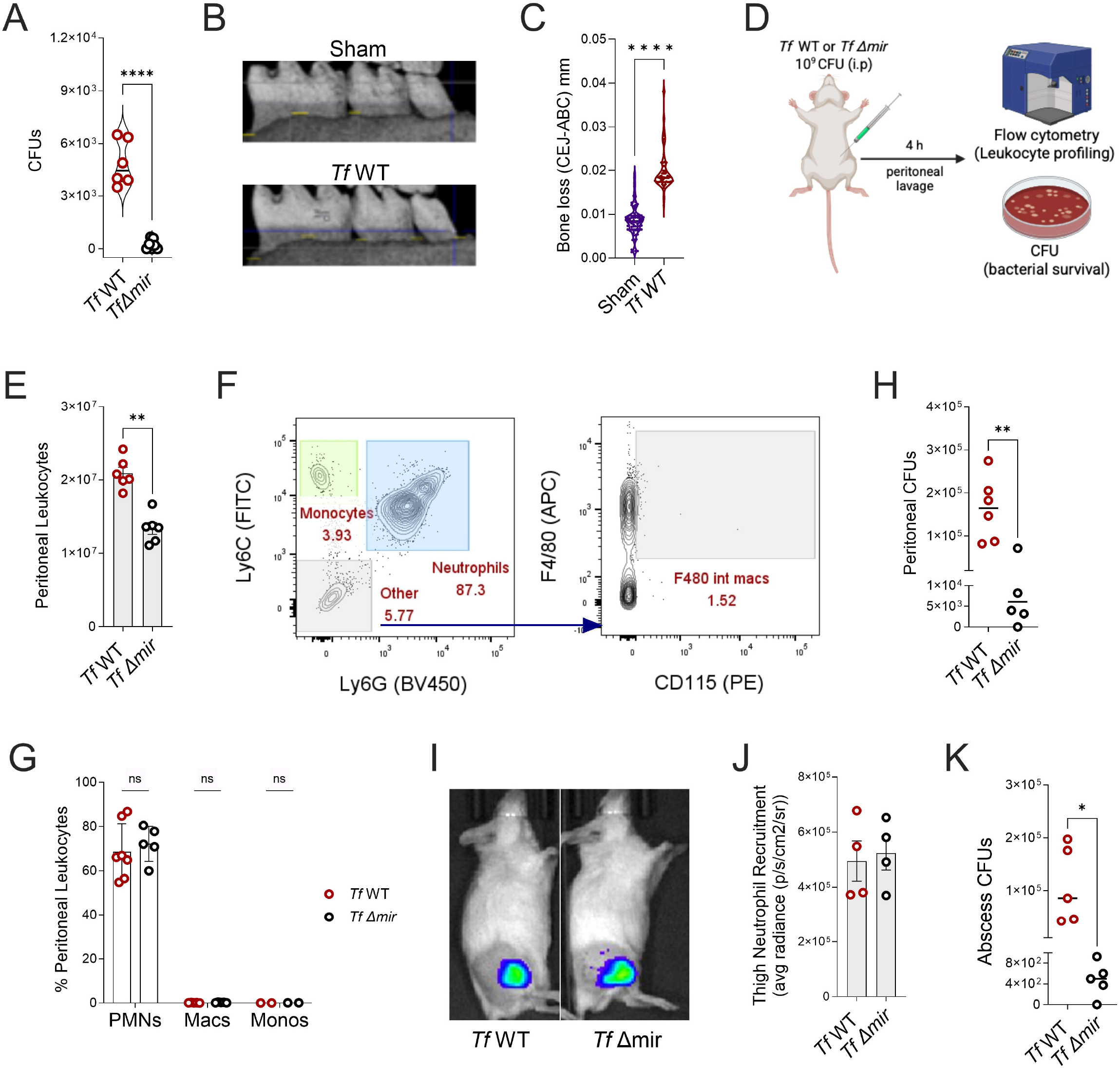
Miropin is essential for *T. forsythia* survival in vivo: **A)** Bacterial (colony-forming units (CFUs)) recovered after 15 min of infection from human neutrophils. Data (mean ± SD) from 6 independent donors are shown. Mice were orally colonized with wildtype *T. forsythia* (WT-*Tf*) for 4 weeks and alveolar bone loss assessed by micro-computer tomography analysis. **B)** shows representative scanned images. **C)** Bone loss/recession was calculated as the distance between the cementoenamel junction and the alveolar bone crest, expressed in mm. **D)** Bacterial peritonitis: schematic of the experimental design. Briefly, mice were intraperitoneally inoculated with 10^9^ CFUs of WT *Tf* or miropin-deleted congenic strain (*Tf*Δ*mir)* for 4 h and peritoneal leukocytes analyzed by flow cytometry. **E)** Total leukocyte counts; **F)** Gating strategy for identification of peritoneal leukocytes based on lineage markers; **G)** Relative % of neutrophils (Ly6G^hi^ Ly6C^int^); monocytes (Ly6C^hi^ CD115^+^ Ly6G^-^); and macrophages (F4/80^+^ CD115^+^); **H)** Surviving bacteria CFUs/ mL in lavage fluid. **I)** Thigh abscess model: Mice were subcutaneously injected with 10^9^ CFUs of WT *Tf* or *Tf*Δ*mir*. 24 h post-bacterium inoculation, neutrophil recruitment (luminol chemiluminescence) was imaged in the IVIS Spectrum imaging system. **I, J)** Representative images and quantification of luminol signal (average radiance) from the infected regions. **K)** CFU/mL of bacteria recovered from abscesses after 24 h. Data are expressed as mean ± SD, and statistical differences for E, H, J and K were determined using a Mann-Whitney t-test or two-way ANOVA with Šídák’s correction (*P<0.05, **P<0.01, ****P<0.0001).

Introducing *Tf* into the peritoneal cavities of mice led to rapid leukocyte infiltration within 4 h, with significantly higher numbers observed in mice challenged with *Tf* WT than in those challenged with *TfΔmir* (**Figure 1D, E**). Flow cytometry analysis of peritoneal lavage showed similar relative numbers of neutrophils, which comprised ∼90% of inflammatory cells in both *Tf* WT- and *Tf-Δmir*-injected mice, with very low levels of macrophages and monocytes (**Figure 1F, G**). *TfΔmir* survival was significantly impaired in the neutrophilic peritoneal environment compared with the *Tf* WT strain (**Figure 1H**). To validate these findings, we used a complemented *Tf* strain with comparable expression of cell membrane glycoproteins TsfA and TsfB within the S-layer and miropin as the WT strain (**Suppl. Figure S1A**). Complementation restored *Tf* survival in the peritoneal cavity (**Suppl. Figure S1B**). We further validated our findings in the thigh abscess model, where injection of *Tf* WT and *Tf*Δmir into the inner thighs of mice led to similar neutrophil recruitment and abscess formation in live mice 24 h post-challenge (**Figure 1I, J**). However, as in the peritonitis model, *Tf*Δmir was cleared quickly, whereas the *Tf* WT strain persisted in abscess exudates (**Figure 1K**). Thus, our *in vivo* and *in vitro* data show that miropin is not only critical for oral colonization but is also essential for *T. forsythia* to resist neutrophil-mediated clearance.

### Miropin does not affect degranulation but blocks the neutrophil oxidative burst

To mechanistically identify the specific antimicrobial effector pathway targeted by miropin, we first expressed and purified wild-type miropin (rMir^WT^) as a recombinant protein, as previously described ^14^, and then assayed neutrophil antimicrobial responses *in vitro*. As a control, activity-dead miropin was generated by replacing the -Val-Lys-Thr-Ser-sequence within the RCL of native or wild-type miropin (rMir^WT^) with Gly-Gly-Gly-Gly-(rMir^G4^), which failed to inhibit purified NSPs (**Suppl. Figure S1C**). In live neutrophils, bacterial killing is mediated by several distinct mechanisms, such as degranulation, activation of the oxidative burst, and formation of neutrophil extracellular traps (NETs) ^2–4^. While miropin did not inhibit the degranulation or mobilization of CD63+ azurophilic granules to the plasma membrane (**Suppl. Figure S1D, E**) in live neutrophils, it potently inhibited the enzymatic activity of NE and CatG released by degranulated (PMA-stimulated) neutrophils in cell-free supernatants (**Figure 2A, B)**. We also tested NE activity during NET release and observed similar inhibition (**Suppl. Figure S1F**).

**Figure 2:**
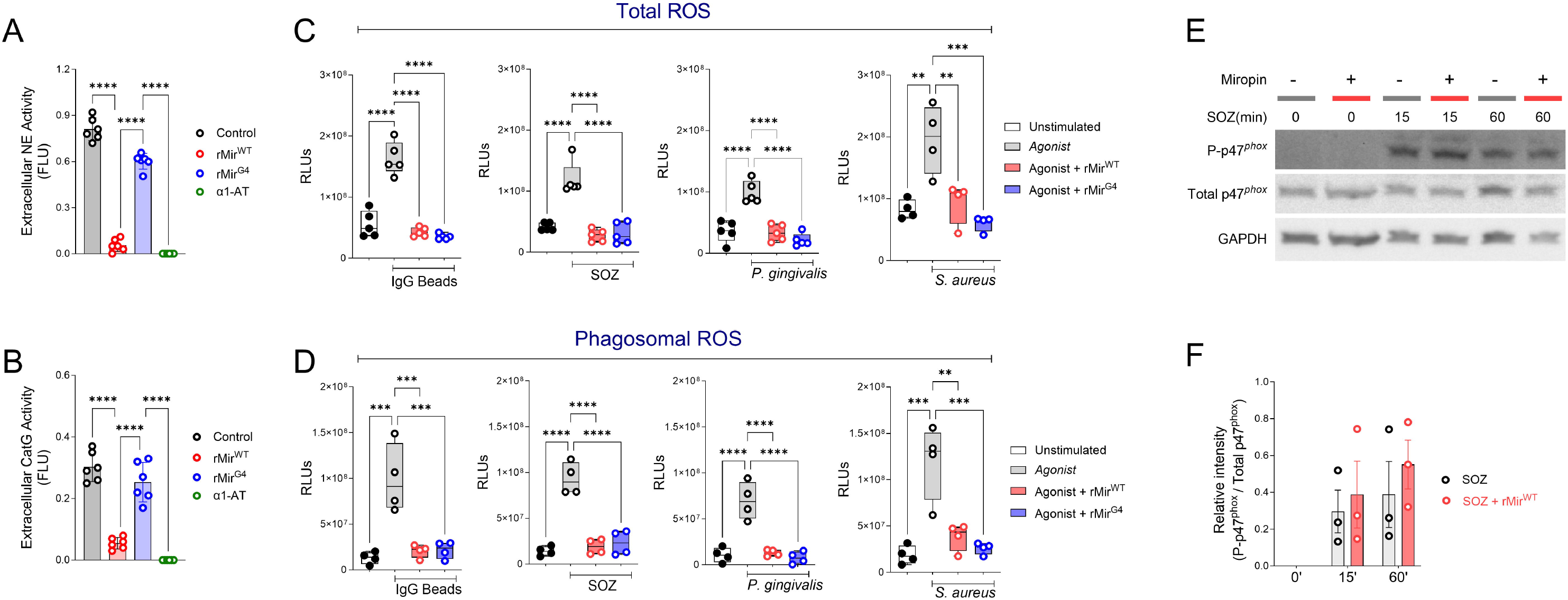
Miropin inhibits neutrophil oxidative burst: A, B) Neutrophils were challenged with 25 nM PMA for 1 h, and **A)** neutrophil elastase and **B)** Cathepsin G activity was determined in cell-free supernatants from 3-4 human donors. **C, D)** Neutrophils were challenged with IgG-coated beads, serum-opsonized zymosan (SOZ), *P. gingivalis*, or *S. aureus* (MOI 100) in the presence or absence of 1 μM rMir^WT^ or rMir^G4,^ and ROS production (total and phagosomal) was measured kinetically as luminol-elicited chemiluminescence over 1 h. Data are shown as integrated relative light units (RLUs) from 3-5 human donors. **E)** phospho-p47*^phox^*(Ser375), total p47*^phox^*, and GAPDH levels were estimated by western blots in human neutrophils stimulated with SOZ in the presence or absence of rMir^WT^. **F)** Densitometry of phospho-p47*^phox^*(Ser375) normalized to p47*^phox^* from 3 independent experiments. Data are represented as mean ± SD and statistical differences determined using one-way ANOVA with Tukey’s multiple comparison test or two-way ANOVA with Šídák’s correction(**P<0.01, ***P<0.001, ****P<0.0001).

Next, we investigated whether miropin confers resistance to the neutrophil oxidative burst, a powerful antimicrobial effector response that operates independently of degranulation, NETosis, and NSP activation. In neutrophils, most ROS generation relies on the activity of the leukocyte NOX2 oxidase, which is assembled on plasma or phagosomal membranes by recruitment of cytosolic subunits during activation. Superoxide (O_2_^-^), the direct product of NOX2 activation, is further converted (dismutated) to H_2_O_2_, which serves as a substrate for MPO-mediated halogenation, generating hypochlorous acid (HOCl), a potent antimicrobial radical species ^2,4^. We measured reactive oxygen species (ROS) generation using a standard luminol-based chemiluminescent assay that depends on MPO-mediated oxidation of luminol to generate light ^23,24^. Stimulation of neutrophils with prototypical particulate agonists of Fcγ receptors (IgG-coated latex beads), complement receptor 3 (serum-opsonized zymosan), or Toll-like receptor (TLR) activating bacterial pathogens *Porphyromonas gingivalis* and *Staphylococcus aureus,* all led to robust ROS generation, which was dramatically and unexpectedly inhibited in the presence of miropin regardless of the variant used (**Figure 2C, D**). We found that miropin did not impact the phosphorylation of the cytosolic p47*^phox^* subunit of the NOX2 oxidase in SOZ-stimulated cells (**Figure 2E, F**) or *P. gingivalis*-challenged neutrophils (**Suppl. Figure S2A, B**), indicating that miropin targeted the upstream MPO-catalyzed step in the ROS amplification cascade. This was further confirmed by the inability of miropin to inhibit ROS in murine peritoneal macrophages, a cell type that lacks MPO (**Suppl. Figure S2C**).

Since rMir^WT^ and rMir^G4^ equally inhibited ROS generation (**Figure 2C, D**), we surmised that miropin-mediated inhibition of the neutrophil oxidative burst occurred independently of the amino acid sequence within the RCL domain. To confirm this, in addition to rMir^WT^ and rMir^G4^, we tested two other recombinant variants of miropin in which the V-K-T motif at the active site of the RCL was substituted with R-V-K and -V-A-T-, which are active serpins with altered inhibitory specificity ^14^. Our data show that miropin variants inhibited MPO regardless of the amino acid sequence within the RCL (**Suppl. Figure S2D, E)**. Collectively, these data indicate that miropin’s serpin function is independent of its MPO-inhibitory capacity and that miropin might moonlight as a peroxidase inhibitor.

### Miropin binds to MPO with high affinity and inhibits its activity

To assess miropin-MPO interactions, we measured the binding affinity between miropin and human MPO using ELISA and isothermal micro-thermophoresis (MST), and found a *K_D_* in the nanomolar range, indicating high binding affinity (**Figure 3A, B).** Next, we performed MPO activity assays and found that miropin potently inhibits MPO activity in a concentration-dependent manner, similar to 4-aminobenzoic acid hydrazide (4-ABH), a synthetic MPO inhibitor (**Figure 3C**). Furthermore, miropin’s capacity to inhibit peroxidases was not limited to a single species, as miropin inhibited human and murine MPO as well as horseradish peroxidase (HRP) (**Figure 3D**). Finally, we tested whether miropin expression was essential for *T. forsythia* to inhibit MPO. We previously showed that miropin is not only secreted into the bacterial culture media but is also exposed on the *T. forsythia* cell surface ^14^, where it could interact with neutrophils. Consistent with this, we found that *Tf-*WT inhibited MPO in a dose-dependent manner, whereas *TfΔmir* did not (**Figure 3E**). Finally, to confirm that miropin inhibits MPO in live neutrophils in response to various stimuli, we used the cell-permeable reporter dye, 3’-(p-Aminophenyl) fluorescein (APF), which is converted to highly fluorescent fluorescein upon interaction with MPO-generated oxidants ^25^. Consistent with the luminol data (**Figure 2C, D**), we observed miropins reduced APF fluorescence (∼50%) in the stimulated live neutrophils, confirming MPO inhibition (**Figure 3F**).

**Figure 3:**
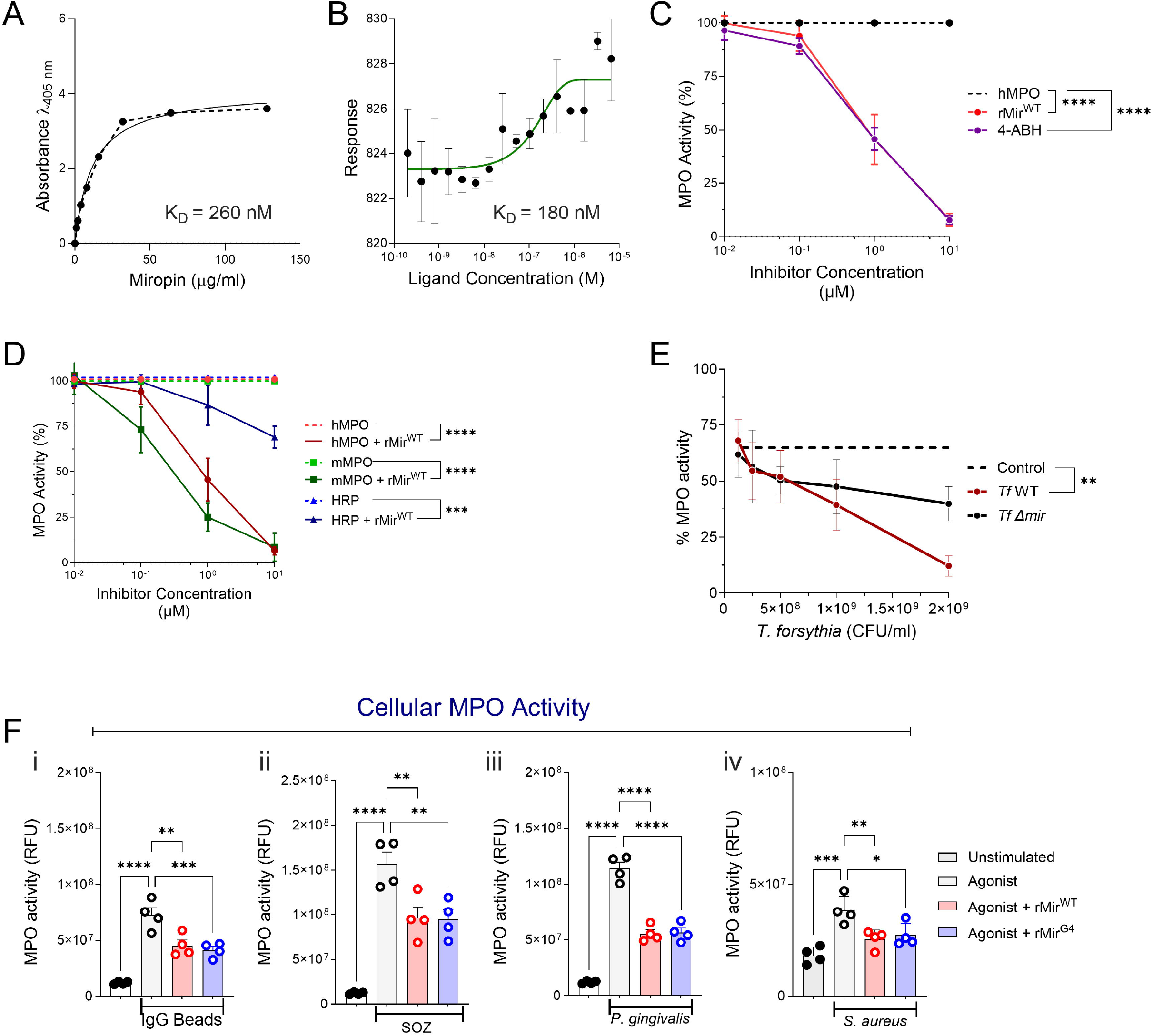
Miropin binds and inhibits myeloperoxidase: **A)** ELISA-based binding assay of rMir^WT^ to myeloperoxidase (MPO). Absorbance was plotted against rMir^WT^ concentration, and a nonlinear regression curve was used to calculate the dissociation constant (K_d_). **B)** Microscale thermophoresis (MST) analysis of the interaction between rMir^WT^ and MPO. Changes in normalized fluorescence (F_norm_) are shown. Data points represent experimental values, and the solid line indicates the nonlinear regression fit used to calculate the K_d_. **C)** Purified human MPO (hMPO) and the synthetic MPO inhibitor 4-ABH; or **D)** peroxidases (hMPO, murine MPO, and horseradish peroxidase [HRP]) were incubated with increasing concentrations of rMir^WT^, and residual MPO activity was determined by oxidation of TMB (3,3’,5,5’-tetramethylbenzidine) over 10 minutes and normalized to MPO alone. **E)** MPO activity was also determined upon incubation with WT *Tf* or *Tf*Δmir cell suspension as described in D and C. For C-E, data are expressed as mean ± SD from three experiments, each performed in triplicate. Statistical differences were determined by one-way ANOVA of the area under the curve (AUC) with Šídák’s correction. **F)** Neutrophils from 3-4 donors were infected with (i) IgG beads, (ii) SOZ, (iii) *P. gingivalis*, or (iv) *S. aureus* (MOI 100) in the presence or absence of rMir^WT^ or rMir^G4^. MPO chlorination activity was measured kinetically over 60 mins by monitoring the increase in APF fluorescence (relative fluorescence units (RFUs)). Data are expressed as mean ± SD, and statistical differences were determined by one-way ANOVA with Dunnett’s multiple comparison test (*P<0.05, **P<0.01, ***P<0.001, ****P<0.0001).

To determine whether miropin inhibits MPO activation within neutrophil phagosomes in a cell-intrinsic manner, we used a flow cytometry-based approach. Neutrophils were challenged with zymosan or bacteria labeled with the acid-sensitive dye pHrodo-succinimidyl ester (pHrodo-red), which fluoresces upon phagocytosis due to acidification ^26^ of the phagosomes, allowing for the specific identification of phagocytosing neutrophils within a population. In the absence of miropins, neutrophils showed a homogeneous population of highly fluorescent APF^hi^ cells, indicating strong MPO activity. However, in the presence of miropins (rMir^WT^ and rMir^G4^), there was considerable heterogeneity, resulting in lower fluorescence and segregation of neutrophils into distinct subsets within the APF^low^ and APF^int^ (intermediate) gates, indicating sluggish or reduced MPO activity in neutrophils challenged with a fungal agonist (zymosan) (**Figure 4A-C**) and *P. gingivalis* (**Figure 4D-F**). We found reduced APF-MFI in neutrophils ingesting WT *Tf* compared to *Tf*Δmir (**Figure 4G-I**), confirming that miropin, which is surface-expressed on *Tf* membranes ^14^, can inhibit MPO within neutrophil phagosomes.

**Figure 4:**
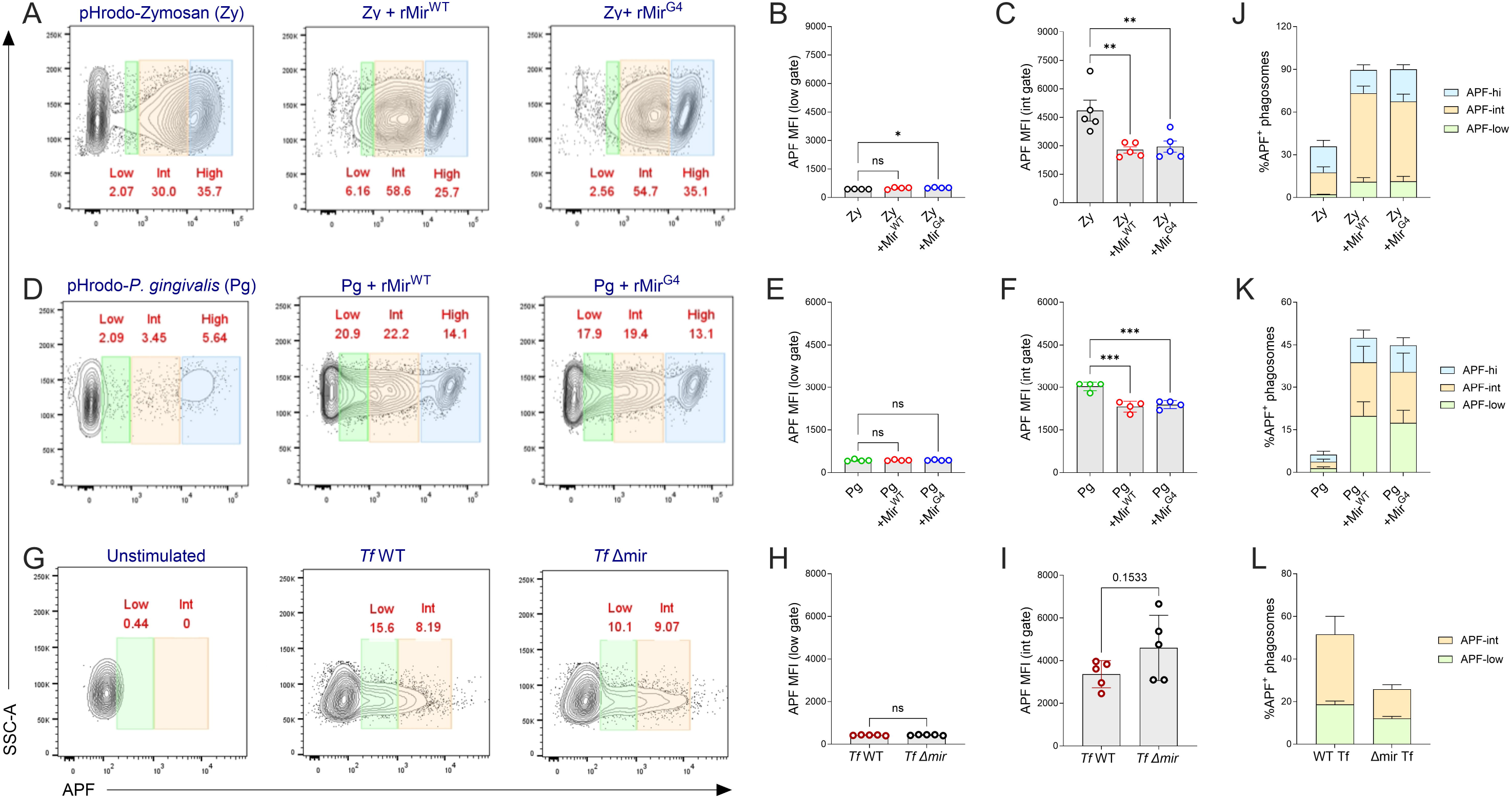
Miropin inhibits MPO within neutrophil phagosomes: Human neutrophils were challenged with pHrodo-labeled phagocytic cargo (zymosan, *P. gingivalis*, *T. forsythia*) for 30 mins in the presence or absence of 1 µM rMir^WT^ or rMir^G4,^ and APF fluorescence was quantified in phagocytosing (pHrodo-positive) neutrophils by flow cytometry. **A-C)** Representative dot plots and quantification of APF mean fluorescent intensity (MFI) within APF^low^ and APF^int^ gates for neutrophils challenged with pHrodo-zymosan. **D-F)** Representative dot plots and quantification of APF mean fluorescent intensity (MFI) within APF^low^ and APF^int^ gates for neutrophils challenged with pHrodo-*P. gingivalis*. **G-I)** Representative dot plots and quantification of APF mean fluorescent intensity (MFI) within APF^low^ and APF^int^ gates for neutrophils challenged with pHrodo-*T. forsythia*. **J-L)** Total phagocytosing (APF positive) neutrophils under each condition. Data are expressed as mean ± SD, and statistical differences were determined using Welch’s t-test or one-way ANOVA with Dunnett’s multiple comparison test (*P<0.05, **P<0.01, ***P<0.001).

### Miropin primes neutrophils via TLR2 for increased phagocytic capacity

In our assays, independent of the nature of the stimulus, neutrophils exposed to miropins seemed to have increased rates of phagocytosis. This was reflected by the increase in the total numbers of pHrodo-positive (**Suppl. Figure S3**) and APF-positive (**Figure 4 J-L**) dual-positive neutrophils. A confounding factor in these observations is that an upward shift in pHrodo MFI (**Suppl. Figure S3**) can result from increased particle uptake (multiple phagosomes per cell) or increased phagosomal acidification, or both. Thus, to determine whether miropin directly affected phagocytosis, we used imaging flow cytometry, which allows simultaneous assessment of pHrodo MFI and particle uptake within the same cell. Spot-counting algorithms were used to identify discrete fluorescent “spots,” or phagosomes, within the boundary of a single cell, represented as subregions, where increased phagocytosis corresponds to an increased number of subregions within a population. Using pHrodo-zymosan as a model particulate agonist, we observed a significantly higher number of spots in miropin-treated cells, with populations segregating into two distinct subregions based on relative particle uptake per cell (**Figure 5A**-**B**). Despite heterogeneity in phagosomal acidification, as reflected by the clustering of cells into pHrodo high, intermediate, and low gates, we did not detect any major changes in acidification in the miropin-treated or control groups (**Figure 5C, D**). Thus, miropin appears to enhance the phagocytic capacity of neutrophils, leading to the uptake of multiple targets. To understand the underlying mechanism, we considered the hypothesis that miropin primes neutrophils, enhancing their phagocytic capacity.

**Figure 5:**
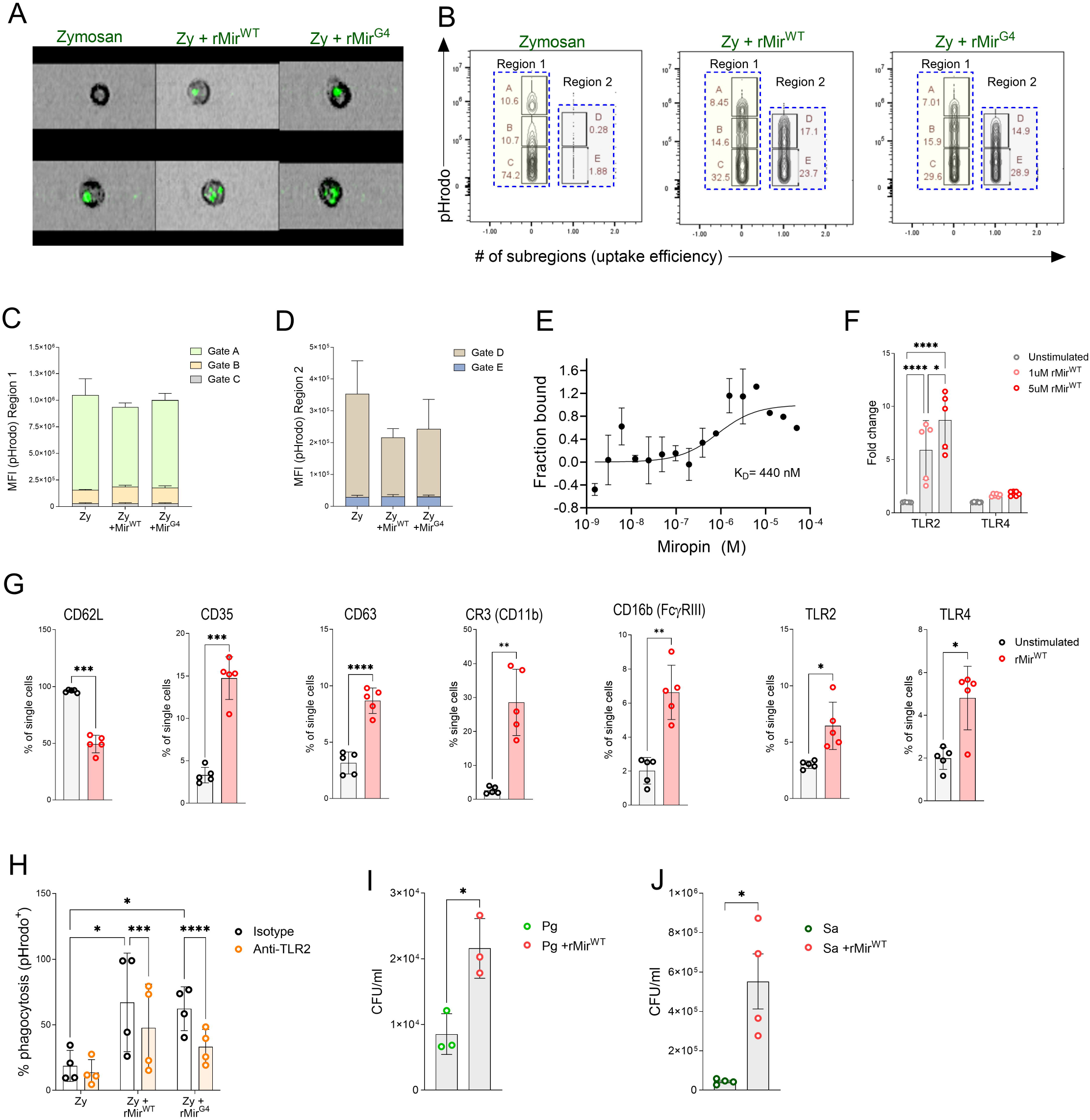
Miropin increases phagocytic uptake in neutrophils: **A)** Representative imaging flow cytometer images of neutrophils challenged with pHrodo-Zymosan in the presence or absence of rMir^WT^ and rMir^G4^ for 30 min. **B)** Bivariate plots showing phagocytic uptake efficiency on the x-axis as # of sub-regions per cell and the y-axis as phagosomal acidification (pHrodo zymosan MFI). Cumulative pHrodo fluorescence (MFI) across all gates in **(C)** region 1 and **(D)** region 2. **E)** Microscale thermophoresis (MST) analysis of the interaction between rMir^WT^ and recombinant human TLR2 (rTLR2). Changes in fraction bound are shown. Data points represent experimental values, and the solid line indicates the nonlinear regression fit used to calculate the K_d_. **F)** HEK-293 TLR2 Blue and HEK-293 TLR4 Blue cells were treated with miropin for 24 h, and SEAP activity was detected spectrophotometrically in supernatants as a measure of TLR activation. **G)** Human neutrophils (4 donors; technical duplicates) were incubated with rMir^WT^ for 1 h and surface receptor expression determined by flow cytometry. **H)** Human neutrophils were pretreated with a neutralizing 5 µg/ml anti-TLR2 antibody or isotype control prior to rMir^WT^ treatment and pHrodo uptake determined by flow cytometry. **I, J)** Neutrophils were infected for 30 min with *P.* gingivalis or *S. aureus* (MOI 1:10) in the presence or absence of rMir^WT^ and rMir^G4,^ and bacterial survival was quantified by enumeration of colony-forming units (CFUs). Data are expressed as mean ± SD, and statistical differences were determined using a Welch’s *t*-test, one-way ANOVA with Tukey’s multiple-comparisons test, or two-way ANOVA with Šídák’s correction (*P<0.05, **P<0.01, ***P<0.001, ****P<0.0001).

Priming is a physiological process in which suboptimal activation of immune receptors enhances neutrophil responsiveness to subsequent stimulation. It is characterized by phenotypic changes such as increased adhesiveness (CD62L shedding), pro-inflammatory gene expression, limited degranulation, and a notable increase in phagocytic capacity via upregulation of phagocytic receptors ^27,28^. We reasoned that miropin signals by binding to TLR2, because it is far more abundant on the surface of human neutrophils than TLR4 ^29^, and previous studies show that *T. forsythia* virulence factors bind and activate TLR2 ^30^. First, we used dot blot assays to confirm reciprocal interactions between miropin and TLR2 by immobilizing either protein on nitrocellulose membranes (**Supp Figure 4B**). Microscale thermophoresis assays showed a K_D_ in the nanomolar range (K_D_ = 440 nM), confirming high-affinity binding between TLR2 and miropin (**Figure 5E**). Next, to link receptor engagement to biological outcomes, we used HEK293 reporter cell lines in which TLR engagement leads to a downstream signaling cascade that ultimately results in the release of secreted embryonic alkaline phosphatase (SEAP) into the cellular supernatant. rMir^WT^, as well as *T. forsythia* strains, preferentially activated TLR2, with minimal responses in a HEK-TLR4 reporter line. Furthermore, loss of miropin, which is immobilized on the membranes of live bacteria, led to reduced TLR2 activation (**Figure 5F and Supp Figure 4A**).

In live neutrophils, rMir incubation led to CD62L shedding and increased surface expression of degranulation markers CD35 and CD63; phagocytic receptors FcγRIII and CR3 (CD11b); and immune receptors TLR2 and TLR4 (**Figure 5G**). Finally, antibody-mediated neutralization of TLR2 reduced the phagocytic capacity of miropin-primed neutrophils, resulting in significantly reduced uptake of pHrodo zymosan in anti-TLR2-treated but not isotype control antibody-treated neutrophils (**Figure 5H**). Interestingly, blockade of TLR2 also prevented miropin-mediated upregulation of zymosan binding receptors such as CD11b, amongst others (**Supp Figure 4C**). Interestingly, despite engagement, we observed only phenotypic, not functional, priming of neutrophils, as miropins effectively block both oxidative and non-oxidative killing in these cells. Thus, despite increased phagocytic capacity, miropin-exposed neutrophils were conducive to bacterial survival (**Figure 5 I, J**).

## DISCUSSION

Homeostatic regulation of proteolysis is essential for controlling several cellular and physiological processes and is achieved in part by proteinaceous inhibitors that block serine, cysteine, metalloproteases, and aspartyl proteases. In vertebrates, these include serpins, TIMPs (tissue inhibitors of metalloproteases), and cystatins (cysteine protease inhibitors). In humans, serpins play an important role in the homeostatic regulation of protease activity by modulating thrombotic and thrombolytic cascades during severe systemic infections, such as sepsis and trauma ^31,32^. While host serpins constrain proteolytic activity and preserve tissue homeostasis, pathogen-derived serpins can exploit similar molecular strategies to circumvent or evade host defenses. Viruses have co-evolved with humans for centuries and are archetypal examples of pathogen-encoded serpins that manipulate the host response. For example, in myxoma viruses, three serpins, SERP1, SERP2, and SERP3, have been identified that target mammalian caspases, thereby blocking apoptosis of the infected cell. SERP-1 can target and modulate thrombotic and complement proteases as a self-defense strategy to evade clearance ^33^. However, serpin distribution in prokaryotes is even more sporadic than in viruses, and their role in bacterial virulence is incompletely understood ^34^.

To date, only a handful of serpins have been described in the human microbiota, mostly among bacteria colonizing the gastrointestinal tract. ‘SERPINBL’ from *Bifidobacterium longum* inhibits pancreatic elastase and NE, forming covalent complexes that were detected in murine fecal pellets. Thus, *B. longum* serpins limit proteolysis in the highly competitive environment of the gut and possibly aid bacterial colonization ^35^. Interestingly, SerpinBL was also shown to inhibit activation of protease-activated receptor 1 (PAR1), thereby attenuating nerve activation and pain in patients with irritable bowel syndrome. A similar observation was made with ‘siropins,’ serpins produced by another human commensal bacterium, *Eubacterium sireaum*, that inhibited NSPs NE and Proteinase 3, possibly suggesting an overlapping role with *B. longum*-derived serpins and a beneficial effect consistent with the commensal nature of these colonizers ^36,37^. In stark contrast, miropin is the first serpin discovered by our group in the oral microbiome ^13^ and is produced by a bacterium long known to contribute to the pathogenesis of periodontitis. It actively contributes to bacterial virulence by priming neutrophils and disengaging distinct neutrophil antimicrobial functions in addition to its serpin function. Thus, miropin functions more as a novel virulence factor by which *T. forsythia* can alter the outcomes of neutrophil-bacterial and interbacterial interactions.

*T. forsythia* colonizes the oral subgingival niche alongside other asaccharolytic periodontal pathogens that are avid protease producers and rely on proteolysis and utilize peptides as a source for nitrogen and carbon. We recently showed that miropin inhibits a protease (lysine-gingipain (Kgp)) produced by its accomplice in periodontitis pathogenesis, *P. gingivalis,* thereby hampering the virulence potential and *in vitro* growth of *P. gingivalis* ^14^. Thus, miropin may have evolved to protect *T. forsythia* surface proteins from proteolytic degradation by both bacterial and host proteases. Consistent with this, we found that miropin inhibits proteases with divergent specificities by using three distinct peptide bonds: Val^367^-Lys^368^, Lys^368^-Thr^369,^ and Thr^369^-Ser^370^ located in the RCL region ^15^. This remarkable plasticity of miropin RCL also enables it to inhibit NSPs ^13^ and plasmin ^38^ proteases that degrade bacterial envelope proteins, thereby inducing bacterial lysis. Unexpectedly, we found that miropin also acts as a potent inhibitor of MPO, independent of its protease-inhibitory activity.

To our knowledge, only three proteins have been characterized as MPO inhibitors so far. These include ceruloplasmin ^39^, staphylococcal peroxidase inhibitor (SPIN) produced by *S. aureus* ^40^, and enterobactin ^41^ produced by *Escherichia coli*. The latter two proteins suppress host innate immune responses, thereby contributing to the virulence of bacterial strains that express them. SPIN proteins are very strong binders of MPO (*K_d_* in the 2-10 nM range) with a well-characterized mechanism of inhibition ^40^. For miropin, binding is weaker (*K_d_* between 180 and 260 nM) but still sufficient to inhibit human MPO activity (EC_50_ in the submicromolar range) and to compromise neutrophil bactericidal activity by blocking MPO-mediated amplification of the neutrophil oxidative burst.

We found that miropin signaled through TLR2 and primed neutrophils to increase their phagocytic capacity, resulting in increased uptake of *P. gingivalis*, *S. aureus,* and the wild-type *T. forsythia* strain. Paradoxically, despite increased phagocytosis, we found that ingested bacteria survived longer within phagosomes in the presence of miropins due to simultaneous inhibition of NSPs and MPO activity. Pathogen engulfment in phagosomes triggers the coordinated maturation and acidification of the phagosome, as well as the activation of lysosomal proteases. Unlike macrophage phagosomes, which undergo progressive acidification, neutrophil phagosomes initially become alkaline before undergoing limited acidification to approximately pH 6. This transient alkalization is closely linked to activation of the Nox2 NADPH oxidase. Nox2-mediated transfer of electrons into the phagosome generates superoxide (O_2_^-^), while compensatory proton flux neutralizes the phagosomal pH ^42^. Therefore, the oxidative burst is more than an isolated antimicrobial mechanism; it shapes the physicochemical environment in which neutrophil granule proteins function, linking the oxidative and non-oxidative arms of neutrophil microbicidal capacity. Furthermore, the neutrophil phagosome is not a uniform compartment, and substantial heterogeneity in phagosomal maturation and antimicrobial activity can occur between neutrophils and within individual cells. Differences in the kinetics of Nox2 assembly, granule fusion, and neutrophil priming or subset state can generate phagosomes with distinct biochemical environments and antimicrobial capacities ^43,44^. Miropin enhanced phagocytic capacity, resulting in multiple phagosomes per cell (**Figure 5**). Although we observed significant heterogeneity in phagosomal acidification (pHrodo fluorescence), we did not observe any consistent differences in phagosomal acidification in the presence or absence of miropin.

Despite TLR2 engagement by miropin and neutrophil priming, we surprisingly did not see any significant changes in inflammatory burden in our in vivo peritonitis and thigh abscess models (**Figure 1**). Neutrophils often feed forward self-amplifying inflammatory loops in vivo via the release of inflammatory cytokines, chemokines, and tissue-degrading proteases that cause tissue damage. NE, Cat G, upon release from degranulating activated neutrophils, degrades elastin, collagen, and other extracellular matrix proteins, while myeloperoxidase activity has been associated with lipid peroxidation and oxidative damage in tissues. Since miropins potently inhibited NSPs and MPO, we think that this would lead to a more controlled inflammatory response despite a higher microbial burden or other antagonizing stimuli. Altogether, our findings expand our understanding of microbial serpins as multifunctional modulators of host defense. Specifically, we show that miropin, the first of its kind serpin in the oral cavity, not only inhibits NSPs but also moonlights as a potent MPO inhibitory protein. This dual functionality could indeed be from distinct protein domains. This ability to simultaneously disengage multiple neutrophil effector responses suggests that miropin functions not simply as a protease inhibitor but as a sophisticated regulator of host antimicrobial responses in the oral cavity, potentially enabling pathogens to persist within an activated host microenvironment while limiting the collateral damage associated with an otherwise robust neutrophil response.

## ACKNOWLEDGEMENTS

These studies were funded by grants DE030939 to JB and JP; 5T32DE032994 to SCM from the National Institute of Dental and Craniofacial Research (NIDCR). The authors thank the Abagail Wexner Research Institute at Nationwide Children’s Hospital Flow Cytometry and Immune Monitoring Core Facility (RRID:SCR_028319) for assistance with cell sorting and data acquisition, respectively. We are grateful to Dr. David Rose for their technical expertise and guidance. The authors also thank Katarzyna Mikruta and Dr. Amir Aliramezani (Department of Microbiology, WBBiB, UJ) for assistance in determining the Kd for the miropin-MPO interaction.

## AUTHOR CONTRIBUTIONS

SCM, FV, DM, MM, MBM, JF, BP, and AF performed the experiments and curated and analyzed data. KZ, BS, MNA analyzed RNA-seq data. SCM interpreted data, made figures and wrote the edited the manuscript with inputs from JP and JB. JP and JB directed all experiments, interpreted data, and wrote the manuscript.

