## Supplemental figures and methods for "A moonlighting serpin of *Tannerella forsythia* compromises the bactericidal activity of neutrophils"

**Supplemental Figure 1**

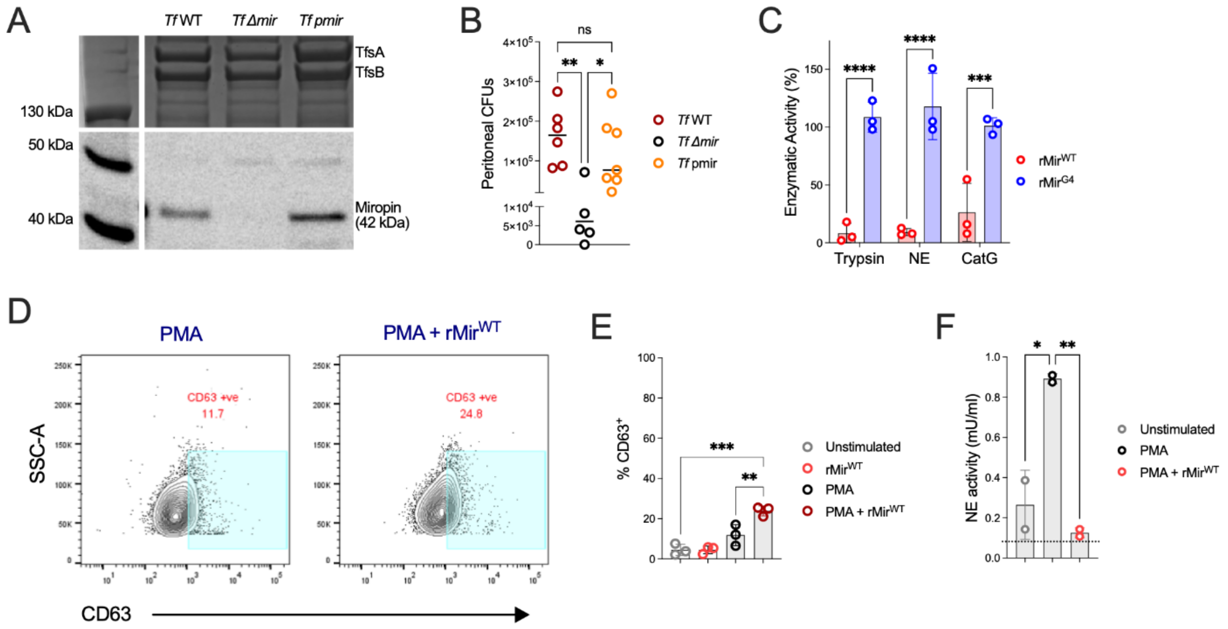


**Supp Figure S1: A)** SDS-PAGE gel showing the expression of S-layer proteins TsfA and TsfB, and Western blot showing expression of miropin in *Tf* wildtype (WT), miropin-deleted (Δmir), and complemented (pmir) strains. **B)** Mice were intraperitoneally challenged with $10^9$ CFUs of WT *Tf*, T*f*Δ*mir*, or *Tf* pmir. After 4 h, peritoneal cavities were lavaged, and bacterial counts (CFU/ mL) in the lavage fluid were enumerated. **C)** Trypsin, neutrophil elastase [NE], and cathepsin G [CatG] were incubated with a 10-fold molar excess of rMir$^{WT}$ or rMir$^{G4,}$ and enzymatic activity was normalized to that of the protease alone. 1 μM rMir$^{WT}$ and rMir$^{G4}$ were estimated as described in the methods. Mobilization of CD63$^{+}$ azurophilic granules in neutrophils stimulated with PMA (20nM) in the presence or absence of 1 μM miropin for 5 min was assessed by flow cytometry. **D)** Representative flow plots; **E)** CD63 expression in neutrophils from 3 human donors.  **F)** NETosis assay looking at the activity of NE on NETs released from PMA-stimulated (50 nM) neutrophils for 4 h.  Data are expressed as mean ± SD and statistical differences were determined using one-way ANOVA with Šídák's multiple-comparisons test or Tukey's multiple comparisons test (*P<0.05, **P<0.01, **P<0.001, ****P<0.0001).

**Supplemental Figure 2**

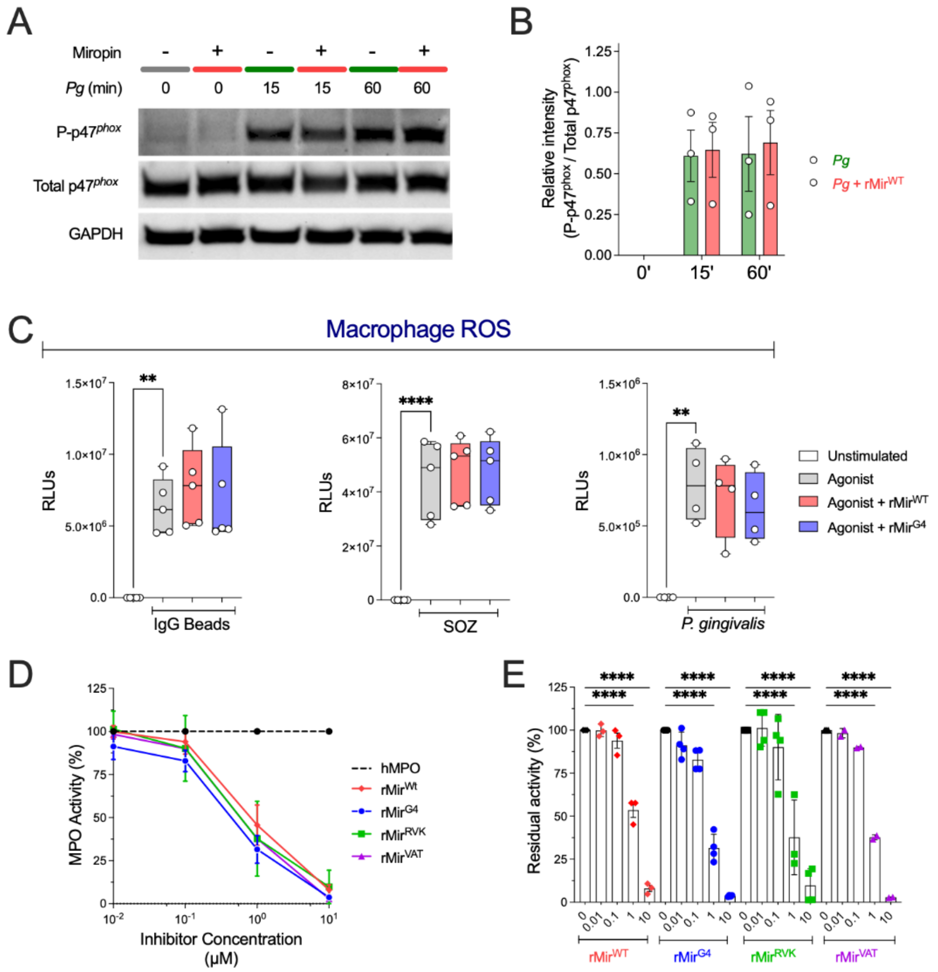


**Suppl. Figure S2: A, B)** Western blots of human neutrophils (3 donors) challenged with *P. gingivalis* (MOI 10) for 15 or 60 minutes in the presence or absence of rMir$^{WT}$, looking at phospho-p47$^{phox}$ (Ser375), p47$^{phox}$, and GAPDH. Densitometry was determined using ImageJ, and phospho-p47$^{phox}$ (Ser375) was normalized to p47$^{phox}$. **C)** Macrophages in the presence or absence of rMir$^{WT}$ were challenged with (i) IgG1-coated beads, or (ii) serum-opsonized zymosan (SOZ) or P. gingivalis (MOI 100), and ROS production was measured kinetically as lucigenin-elicited chemiluminescence. Integrated relative light units (RLU) over 60 min are shown. Data are expressed as mean ± SD and statistical differences were determined using one-way ANOVA with Tukey's multiple-comparisons test (*P<0.05, ***P<0.001, ****P<0.0001). **D)** hMPO was incubated with increasing concentrations of rMir$^{WT}$, rMir$^{G4}$, rMir$^{RVK}$, and rMir$^{VAT}$. Residual MPO activity was determined by TMB oxidation over 10 min and normalized to MPO alone. **E)** Quantification of MPO activity by area under the curve (AUC). Data are shown as mean ± SEM from three independent experiments. Statistical differences were assessed by one-way ANOVA with Šídák's multiple-comparisons test (**P<0.01, ****P<0.0001).

### Supplemental Figure 3

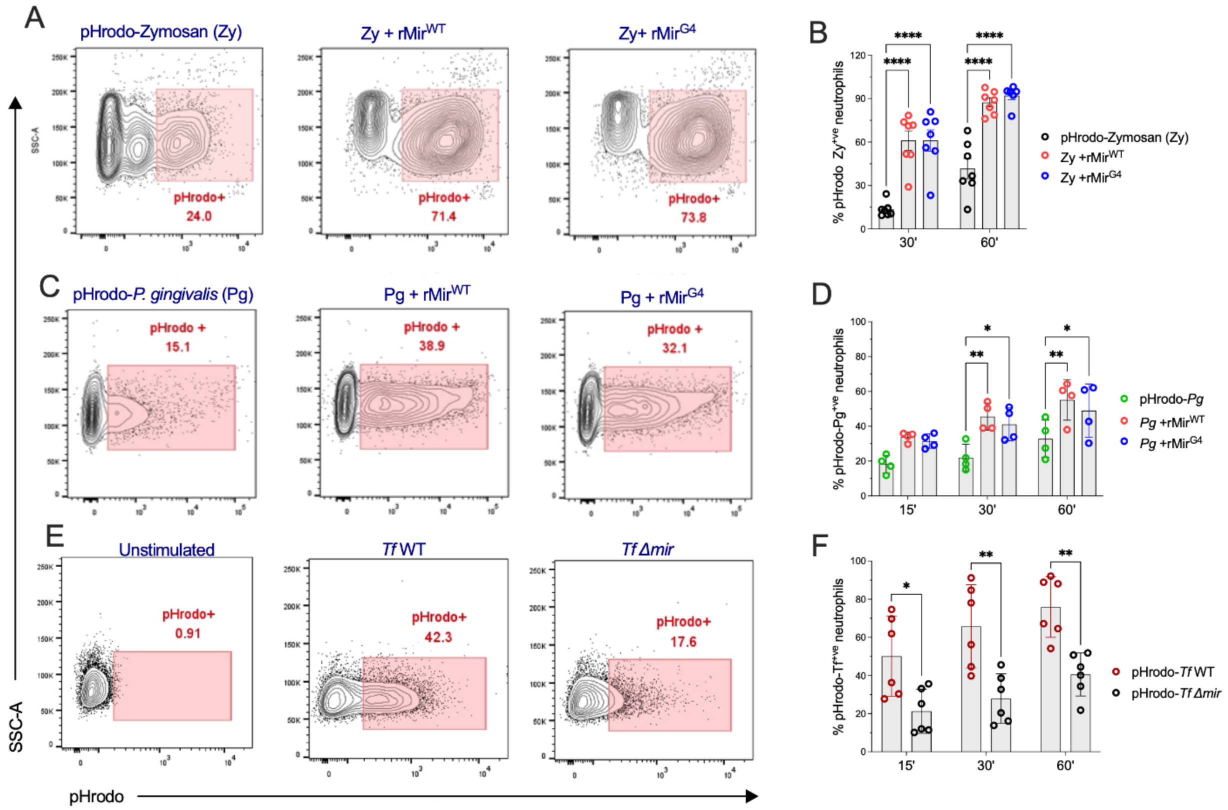


**Supp Figure S3**: Human neutrophils (4-7 donors) were stimulated with **A, B)** pHrodo-Zymosan, **C, D)** pHrodo-*P. gingivalis*, or **E, F)** pHrodo-*Tf* in the presence or absence of $rMir^{WT}$ and $rMir^{G4}$ for various time points. Quantification of pHrodo-positive neutrophils at 15, 30, or 60 min of stimulation from all donors is shown in B, D, and F, respectively. Data are expressed as mean ± SD and statistical differences were determined using two-way ANOVA with Tukey's or Šídák's multiple-comparisons test (*P<0.05, **P<0.01, ****P<0.0001).

### Supplemental Figure 4

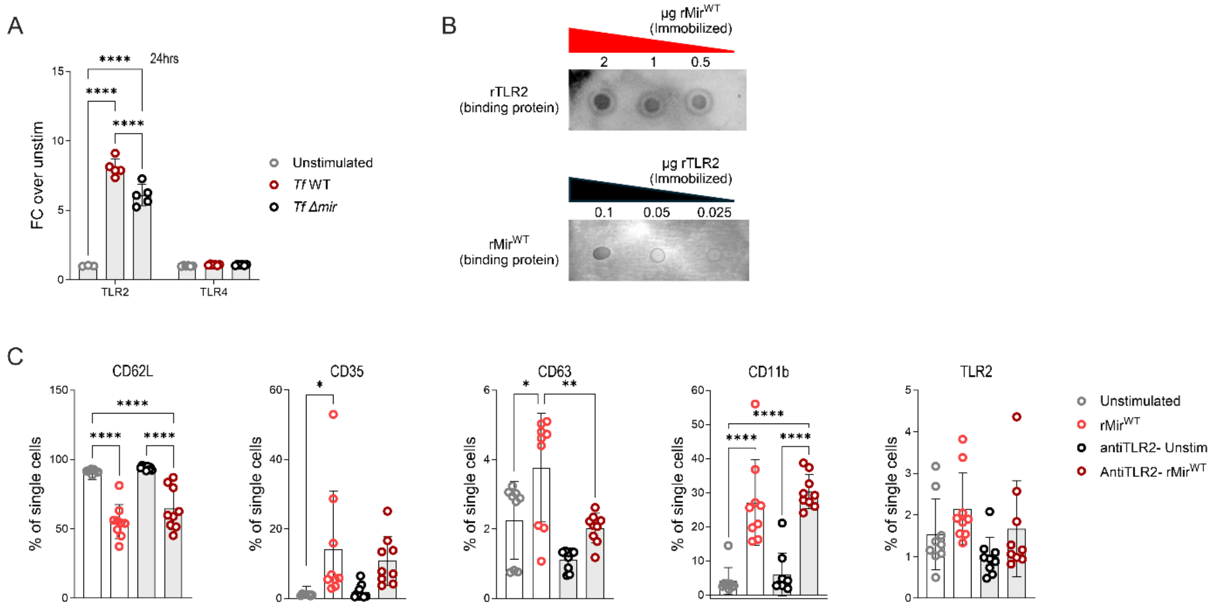


**Suppl Figure S4: A)** HEK293-BLUE reporter cell lines specifically expressing either TLR2 or TLR4 were challenged with *T. forsythia* (WT *Tf* or *TfΔmir*) at MOI 10 for 24 h; TLR activation was determined by the detection of secreted SEAP in the culture supernatant. **B)** Dot blot protein binding assay. Various amounts (2-0.5 μg rMir$^{WT}$ or 0.1-0.025 μg rTLR2) were loaded onto nitrocellulose. Nitrocellulose membranes were then incubated with 1 μg/ml rTLR2 or 2 μg/ml rMir$^{WT}$. **C)** Human neutrophils were pretreated with 5 μg/ml anti-TLR2 or isotype control antibodies prior to 1 h stimulation with rMir$^{WT}$, and the expression of various receptors on the plasma membrane was detected by flow cytometry. Data from 3 donors, run in technical triplicate, are shown. Data are expressed as mean ± SD, and statistical differences were determined using two-way ANOVA with Tukey's multiple-comparisons test (*P<0.05, **P<0.01, ***P<0.001, ****P<0.0001).

### SUPPLEMENTAL METHODS

#### Generation of the *T. forsythia* miropin deletion mutant (Δ*mir Tf*):

A suicide plasmid designed to replace the miropin coding sequence with a chloramphenicol resistance cassette (*cat*) was commercially synthesized [1]. The construct contained 728 bp of upstream and 786 bp of downstream homologous regions flanking the miropin locus of WT *Tf*, separated by the 660 bp *cat* coding sequence. All elements were cloned into the pUC57 backbone to generate plasmid pΔMiropin [1]. pΔMiropin was introduced into electrocompetent WT *Tf* cells by electroporation, and transformants were selected on agar supplemented with chloramphenicol (10 µg/ml). Correct integration of the deletion cassette was confirmed by DNA sequencing.

#### Generation, expression, purification of recombinant miropin variant proteins:

Recombinant miropin (rMir$^{WT}$) and several reactive-center loop (RCL) mutant variants rMir$^{GGGG}$ or (rMir$^{G4}$), rMir$^{RVK}$, and rMir$^{VAT}$ were generated using plasmid pGEX-6P-1, which contains cloned miropin as a template [1]. Site-directed mutagenesis of the $^{367}$VKTS$^{370}$ motif was performed using the Quik-Change Lightning Site-Directed Mutagenesis Kit (Agilent Technologies) according to the manufacturer's instructions. Mutagenic primers were synthesized by Genomed (Warsaw, Poland) using the following sequences:

- WT-mirF; 5' CGTAGAAATGGTAGCAACGTCATCCCCCTC 3'
- WT-mirR: 5' GAGGGGGATGACGTTGCTACCATTTCTACGG 3'
- G$^4$-mirF: 5' GGTGGCGGTGGCTCCCCCTCTACCACTCCTATCAA 3'
- G$^4$-mirR: 5' GGAGCCACCGCCACCCATTTCTACGGCTGTTACGG 3'
- RVK-mirF: 5' GCCGTAGAAATGCGCGTGAAATCATCCCCCTCTACC 3'
- RVK-mirR: 5' GGTAGAGGGGGATGATTTCACGCGCATTTCTACGGC 3'
- VAT-mirF: 5' CCGTAGAAATGGTAGCAACGTCATCCCCCTC 3'
- VAT-mirR: 5' GAGGGGGATGACGTTGCTACCATTTCTACGG 3'

The resulting mutant constructs replaced the native VKTS motif with sequences that alter serpin activity: G4, VAT, and RVK. All constructs were verified by Sanger sequencing. Recombinant miropin proteins were expressed in *Escherichia coli* (Rosetta DE3 cells) and purified by glutathione-Sepharose affinity chromatography followed by size-exclusion chromatography on a HiLoad 16/60 Superdex 75 pg column. The purity of the recombinant proteins was confirmed by SDS-PAGE, and the level of contaminating endotoxin was measured using commercially available endotoxin kits (Thermo-Fisher).

***Tannerella forsythia* identification SDS-PAGE and Western blot:**

*Tannerella forsythia* (WT, Δ*mir*, and pmir) strains were grown as described above. Bacterial cultures were normalized to approximately 1 × $10^9$ CFU and collected by centrifugation at 4,400 rpm for 15 min at room temperature. Cell pellets were washed once with 1× PBS and resuspended in 1 mL PBS. Aliquots (65 µL) were mixed with 4× NuPAGE LDS Sample Buffer and 10× NuPAGE Sample Reducing Agent (Thermo Fisher Scientific) to a final volume of 100 µL. Samples were heated at 100°C for 10–15 min and briefly centrifuged before electrophoresis. Samples were separated on 4–12% Bis-Tris NuPAGE gradient gels (Thermo Fisher Scientific) and either stained with Coomassie Brilliant Blue for visualization of S-layer proteins (TsfA and TsfB) or transferred to membranes for Western blot analysis.

For Western blot analysis, membranes were transferred to a PVDF membrane (Invitrogen) using the Invitrogen Power Blotter for 7 minutes at 120 V. Membranes were blocked in 5% (w/v) skim milk in TBST (20 mM Tris, 0.5 mM NaCl, 0.1% Tween-20; pH 7.5) and sequentially probed with antibodies against probed with a rabbit anti-miropin primary antibody (1:1,000) for 2 h, followed by an HRP-conjugated anti-rabbit secondary antibody (1:1,000) for 1 h. Protein bands were detected using Pierce ECL substrate (Thermo Fisher Scientific) and imaged using a ChemiDoc MP Imaging System (Bio-Rad). Densitometric analysis was performed using ImageJ v1.53e.

**MPO-Miropin binding affinity assays:**

The dissociation constant ($K_D$) between recombinant miropin ($rMir^{WT}$) and purified human MPO (Atlanta Biologicals) was assessed by ELISA and microscale thermophoresis (MST). For ELISA, high-binding 96-well ELISA plates (Thermo Scientific Immuno Clear Standard Modules) were coated with MPO (4 µg/ml in PBS, 100 µl/well) overnight at 4 °C and then blocked with assay buffer (2% bovine serum albumin in PBS) for 1 h at 37 °C. Wells were washed four times with PBS containing 0.05% Tween-20 (TBST). Serial dilutions of $rMir^{WT}$ (1-16 µg/ml) prepared in PBS were added to wells (100 µl/well), and plates were incubated for 1 h at 37 ºC. Following four washes with PBST, bound miropin was detected using 2 µg/ml rabbit anti-miropin primary antibody for 1 h at 37 ºC followed by alkaline phosphatase-conjugated goat anti-rabbit IgG secondary antibody (1:5,000 dilution) for 1 h at 37 ºC. Wells were developed using *p*-nitrophenyl phosphate (PNPP; Thermo Scientific) substrate prepared in 1x diethanolamine buffer according to the manufacturer's instructions (Thermo Scientific). Plates were incubated at room temperature for 10 minutes before reactions were stopped with 1 N NaOH. Absorbance was measured at λ = 405 nm using a Sunrise microplate reader (Tecan, Männedorf, Switzerland).

For MPO-MST assays, a Monolith NT.115 instrument (NanoTemper Technologies GmbH, Munich, Germany) was used. MPO was fluorescently labeled on lysine residues with NT-647 NHS dye and incubated with $rMir^{WT}$ at concentrations ranging from 0.5 nM to 10 μM for 5 min at 20°C. Experiments were performed in PBS (pH 7.4) at 25 °C. Samples were loaded into Monolith NT.115 standard-treated glass capillaries, and initial fluorescence measurements were followed by thermophoresis measurements using 40% LED power and medium MST power. $K_D$ values were calculated using Affinity Analysis software, assuming a 1:1 binding model and using the signal from the default MST-on time.

**TLR2-Miropin binding affinity assays:**

For MST assays, a Monolith NT.115 instrument (NanoTemper Technologies GmbH, Munich, Germany) was used. rTLR2 (R&D Systems, cat. no. 2616-TR) was fluorescently labelled using the His-Tag Labeling Kit RED-tris-NTA 2nd Generation (NanoTemper Technologies) and incubated with miropin at varying concentrations. Experiments were performed in PBS (pH 7.4) at 25°C. Samples were loaded into Monolith NT.115 standard-treated glass capillaries, and initial fluorescence measurements were followed bhoresis measurements using 40% LED power and medium MST power. Data from three independently pipetted measurements were analysed using MO.Affinity Analysis software (NanoTemper Technologies) to determine the apparent $K_D$.

**NADPH oxidase and downstream peroxidase activity assays:**

Reactive oxygen species (ROS) production by human neutrophils was measured kinetically using luminol- or lucigenin-enhanced chemiluminescence as previously described [2,3]. Freshly isolated neutrophils were plated at 2 x $10^5$ cells per well in white 96-well plates. Neutrophils were incubated with 50 μM luminol (Sigma) and 2 U/ml horseradish peroxidase (HRP; Sigma) or 175 µM lucigenin (Cayman Chemicals). Where indicated, extracellular ROS was quenched with 75 U/ml superoxide dismutase (SOD; Sigma). Neutrophils were stimulated with *Pg*, *Tf*, *S. aureus*, or IgG-opsonized latex beads (100:1 particle: cell), or serum-opsonized zymosan (SOZ; 200 μg/ml). Chemiluminescence was recorded continuously for 60 min at 37 ºC using a SpectraMax L microplate luminometer (Molecular Devices), and total integrated relative light units (RLUs) were quantified [4,5].

**MPO activity assays:**

All reactions were performed in 5 mM sodium acetate buffer (pH 5.4), at a final volume of 50 μl. Human MPO (3.57 nM), mouse MPO (6.15 nM), or horseradish peroxidase (HRP) (7 pM) were preincubated with varying concentrations of miropin (0.1 -10 μM), *Tf* ($OD_{600}$ = 1.0) or the MPO synthetic inhibitor 4-aminobenzoic acid hydrazine (4-ABH) for 15 min at room temperature. The substrate solution containing 2.8 mM 3,3',5,5'-tetramethylbenzidine (TMB) and 0.0022% hydrogen peroxide was then added. Peroxidase activity was monitored kinetically by measuring absorbance at λ = 650 nm over 10 min using a BioTek Synergy H1 (Agilent). Activity was calculated from the change in absorbance and normalized to active enzyme concentration using the calculation below:

Enzyme Activity (μmoles/min/ml) =

$$\frac{\Delta A_{650}}{39000M^{-1}cm^{-1} \times pathlength *} \times \frac{total\ volume}{volume\ of\ sample\ (inhibitor\ or\ bacteria)} \times sample\ dilution$$

For plate-based assessment of MPO activity in activated neutrophils, we used 3'-(p-aminophenyl) fluorescein (APF; Thermo Scientific). Freshly isolated neutrophils ($3x10^6$ cells/ml) were suspended in PBS containing 7.5 mM glucose, 0.5 mM $MgCl_2$, and 0.9 mM $CaCl_2$ (PBSG) and incubated with 2 μM APF for 30 minutes at 37 ºC with end-over-end rotation. Cells were challenged with 1 μM $rMir^{WT}$, $rMir^{G4}$, or 4-ABH, then stimulated with IgG-coated latex beads, SOZ, *Pg*, *Tf*, or *S. aureus*, ~~or PMA~~ (as described for NADPH oxidase and peroxidase activity assays). Oxidation of APF to fluorescein was monitored kinetically for 60 minutes at 37 ºC using fluorescence measurements at Ex/Em 490/515 nm. Total integrated fluorescence units (FLUs) were quantified.

**pHrodo Red labeling of bacteria**

Bacteria were harvested by centrifugation at 10,000 x g for 10 minutes and resuspended in 1 mL PBS to an $OD_{600}$ of 5. Then pHrodo red succinimidyl ester (Invitrogen) (1.25 μl of 1.7 mg/ml solution) was added to 1 ml of the bacterial suspension, and the mixture was incubated for 30 minutes at 37 ºC with constant shaking (100 rpm). Bacterial cells were washed three times with PBS. Labeling efficiency was determined by flow cytometry.

**Flow cytometry:**

Flow cytometry data were acquired using the BD LSR Fortessa flow cytometer (BD Biosciences). For imaging cytometry the S8 Cell Sorter (BD Biosciences) was used. Unless otherwise stated, all data were analyzed using FlowJo v10 and v11 software (BD Biosciences/Waters).

**Degranulation assays:**

Freshly isolated neutrophils ($10^6$ cells) were resuspended in RPMI and pretreated with 1 μM $rMir^{WT}$ for 30 min before stimulation with PMA (20 nM) for 5 min at 37 ºC. Reactions were stopped with ice-cold FACS buffer (PBS containing 1% BSA and 2 mM EDTA). buffer. Cells were blocked with 10% human AB serum and stained with antibodies against CD63 (clone H5C6, FITC; Biolegend) for 45 min on ice in the dark. After two sequential washes with ice-cold FACS buffer, relative degranulation was assessed by flow cytometry.

**Neutrophil elastase and cathepsin G activity assay:**

Neutrophils ($1 \times 10^6$ cells/mL) were incubated with 1 μM $rMir^{WT}$, $rMir^{G4}$, or α1-antitrypsin (α1-AT) in the presence or absence of PMA (25 nM). After 1 h, 100 µL of the cell suspension was added to 100 µL of fluorogenic substrate (50 µM AAPV-AMC for neutrophil elastase or AAPF-AMC for cathepsin G) prepared in elastase activity buffer. Fluorescence was monitored at 37°C using excitation and emission wavelengths of 350 and 460 nm, respectively.

**NET-associated neutrophil elastase assay:**

NETosis and NE activity assays were performed according to the manufacturer's protocol (Abcam). Briefly, neutrophils ($1x10^6$) were suspended in NET assay buffer (RPMI, 5% BSA, 1 μM $CaCl_2$) and stimulated with PMA (20 nM) for 4 h to induce NET formation. Following stimulation, wells were gently washed twice with 1 mL pre-warmed NET Assay Buffer to remove soluble neutrophil elastase. NETs were subsequently digested by incubation with S7 Nuclease (15 U/ml) for 15 min at 37°C. Supernatants were collected, and nuclease activity was terminated by addition of EDTA, followed by centrifugation at 300 × g for 5 min to remove cellular debris. NET-associated neutrophil elastase activity in the resulting supernatants was quantified using the Neutrophil Elastase Activity Assay included in the NETosis Assay Kit (Abcam, ab235979). Samples and elastase standards were assayed in duplicate by incubation with the neutrophil elastase substrate for 1–2 h at 37°C. Absorbance was measured at 405 nm, and elastase activity was calculated from a quadratic standard curve after subtraction of the blank.

**NOX2 western blotting:**

To prevent proteolysis of NADPH oxidase subunits, neutrophils were incubated with 2.7 mM diisopropyl fluorophosphate (DFP; Sigma) for 20 min at 4 ºC to inhibit endogenous serine protease activity [6]. After DFP treatment, neutrophils were pre-incubated with 1 µM $rMir^{WT}$ or $rMir^{G4}$ for 30 min, then stimulated with 5 mg/ml serum-opsonized zymosan (SOZ) or *Pg* (10 bacteria: 1 cell) for 15 or 60 min at 37 ºC. At each time point, cells were washed twice with ice-cold PBS.  Cell pellets were suspended in lysis buffer (RIPA buffer (Thermo) containing cOmplete protease inhibitor cocktail (Roche) and PHOSstop phosphatase inhibitor (Roche)), and protein concentration was estimated using the BCA assay. For western blots, samples were treated with NuPAGE sample buffer and reducing agent (Invitrogen) and boiled for 10 min.

Lysates equivalent to $10^6$ neutrophils per well were resolved on 4-12% Bis-Tris gradient gels (Invitrogen) and transferred to a PVDF membrane (Invitrogen) using the Invitrogen Power Blotter for 7 minutes at 120 V. Membranes were blocked in 5% (w/v) skim milk in TBST (20 mM Tris, 0.5 mM NaCl, 0.1% Tween-20; pH 7.5) and sequentially probed with antibodies against phospho-$p47^{phox}$ (Ser345; 1:1000; Invitrogen), total $p47^{phox}$ (1:1,000; Abcam), and GAPDH (1:1,000; Cell Signaling), followed by probing with appropriate HRP-conjugated secondary antibodies (1:1,000 anti-rabbit, Cell Signaling; 1:5,000 anti-goat; Invitrogen). Protein bands were detected using Pierce enhanced chemiluminescence (ECL) substrate (ThermoFisher Scientific) and imaged using ChemiDoc MP (BioRad). Densitometric analysis was performed using ImageJ (1.53e).

**Bacterial survival assay:**

Intracellular bacterial survival within neutrophils was assessed as previously described [3]. Freshly isolated neutrophils suspended in RPMI (no antibiotics) were pre-incubated with 1 µM $rMir^{WT}$ then infected with *P. gingivalis*, *S. aureus*, or *Tf* at an MOI of 10 (10 bacteria: 1 neutrophil). Infections were carried out at 37 ºC for the indicated time points. At each time point, cells were washed twice with PBS and incubated in antibiotic-containing media (300 µg/ml gentamicin and 200 µg/ml metronidazole for *P. gingivalis*; 1 µg/ml flucloxacillin and 25 µg/ml lysostaphin for *S. aureus*) for 30 min at 37 °C. No antibiotics were used for *Tf*. Neutrophils were subsequently washed twice with PBS and lysed in sterile $diH_2O$ for 20 min at room temperature.

Lysates were serially diluted and plated on agar plates for colony-forming unit (CFU) enumeration. All antibiotics were purchased from Millipore-Sigma.

**Oral gavage model:**

Mice were pre-treated with 870 μg/mL sulfamethoxazole and 170 μg/mL trimethoprim for 8 days to deplete oral microflora. On day 10, mice were orally infected with $1 \times 10^{10}$ CFU of *Tf* WT or *TfΔmir* suspended in 2% sterile CMC by oral gavage on alternate days for a total of 2 weeks. Mice were euthanized 4 weeks after the last bacterial inoculation. Alveolar bone loss was assessed by micro-computed tomography (μCT) to measure the distance from the cementoenamel junction (CEJ) to the alveolar bone crest (ABC) [7].

**Murine abscess model and *in vivo* imaging**

Subcutaneous abscess infections were performed as previously described, using B6.Cg-Tyrc-J/J mice [8]. Briefly, injection sites were shaved and depilated 24 hours prior to injection. Mice were inoculated subcutaneously in the outer thigh with 100 μl of *Tf* ($8x10^8$ CFUs) suspended in PBS. Mice infected with *Tf* were euthanized 24 h post-infection. Abscesses were surgically excised, homogenized, serially diluted, and plated for CFU enumeration.

For inflammatory oxidant imaging, mice received intraperitoneal (i.p.) injections of luminol (300 mg/kg), and imaged at 4 h post-infection. Mice were anesthetized with 1-2% isoflurane and imaged using an IVIS Spectrum imaging system (Revvity). Images were acquired for 1 minute at f/stop 1 with binning set to 16. Bioluminescence signals were quantified as radiance (photon/sec/$cm^2$/sr) using Living Image software v4.7.4 (Caliper Life Sciences) [9].

**Murine peritonitis model:**

Experimental bacterial peritonitis was induced by intraperitoneal injection of *Tf* ($10^9$ CFUs) suspended in 1 ml of PBS, as previously described [10]. Four hours post-infection, mice were euthanized, and the peritoneal cavity was lavaged with 8 ml of PBS containing 2 mM EDTA. The recovered lavage fluid was serially diluted and plated for CFU enumeration. Peritoneal exudate cells were treated with Fc receptor blocking reagent (2.4G2 hybridoma supernatant) then stained with antibodies against Ly6G (clone 1A8, BV605; BD Biosciences), Ly6C (clone HK1.4, FITC; Biolegend), F4/80 (clone BM8, APC; Biolegend), and CD115 (clone AFS, PE; BD Biosciences) for 45 min on ice in the dark and analyzed by flow cytometry.

**HEK-Blue TLR2 and TLR4 cultures:**

HEK-Blue TLR2, and HEK-Blue TLR4(NF-κB reporter cell lines (InvivoGen) were maintained in culture in DMEM +10% ΔFBS supplemented with 10ug/ml normocin (Invivogen) and, when applicable, 100 ug/ml zeomycin (Invivogen). For TLR activation assays, cells were treated with 1 or 5 μM $rMir^{WT}$ or challenged with *Tf* at an MOI of 10 for 24 h. TLR activation was assessed by measuring secreted embryonic alkaline phosphatase (SEAP) activity in cell-free supernatants using Quanti-Blue substrate, at $\lambda_{620}$ nM using a BioTek Synergy H1 (Agilent).

**Dot blots for miropin binding:**

Miropin was serially diluted 2-fold and 4-fold in PBS. Nitrocellulose strips (1 × 3 cm) were spotted with 2 μL of each miropin dilution, corresponding to 2, 1, and 0.5 μg of protein, with 1 cm spacing between spots. After air-drying for several minutes, the strips were incubated in blocking solution for 2 h, washed, and incubated with recombinant human TLR2 (rTLR2; 1 μg/mL) for 2 h. Strips were washed three times and incubated with anti-TLR2 antibodies (2 μg/mL) for 2 h, followed by incubation with the appropriate secondary antibodies for 2 h. After three additional washes, the reaction was developed. For the reciprocal binding assay, rTLR2 was serially diluted 2-fold and 4-fold in PBS, corresponding to 0.1, 0.05, and 0.025 μg of protein and spotted onto nitrocellulose strips as above. After air-drying for several minutes, the strips were incubated in blocking solution for 2 h, washed, and incubated with rMir (2 μg/mL) for 2 h. Following three washes, the strips were incubated with anti-miropin antibodies for 2 h, followed by the appropriate secondary antibodies for 2 h. The strips were washed three times before developing the reaction.

### SUPPLEMENTAL REFERENCES (FOR METHODS)


1. Ksiazek, M., Benedyk-Machaczka, M., Sochaj-Grzegorczyk, A., Mizgalska, D., Waligorska, I., Nowak, M., Kaczor, A., Potempa, B., Fuchs, A., Thogersen, I.B., et al. (2025). Innocent until proven guilty: Tannerella forsythia may attenuate the virulence of Porphyromonas gingivalis. bioRxiv. 10.1101/2025.09.05.674453.
2. Cooper, K.N., and Bagaitkar, J. (2020). Altruistic death: Neutrophil apoptosis maintains gingival health. Journal of Leukocyte Biology *108*, 1497-1499. 10.1002/jlb.5ce0520-122r.
3. Cooper, K.N., Wielento, A., Morris, S.C., Terekhova, M., Rodriguez-Hernandez, C.J., Potempa, B., Carey, K.A., Artyomov, M.N., Potempa, J., and Bagaitkar, J. (2025). Limited proteolysis of neutrophil granule proteins by the bacterial protease RgpB depletes neutrophil antimicrobial capacity. Journal of Leukocyte Biology *117*. 10.1093/jleuko/qiae209.
4. Bagaitkar, J., Barbu, E.A., Perez-Zapata, L.J., Austin, A., Huang, G., Pallat, S., and Dinauer, M.C. (2017). PI(3)P-p40phox binding regulates NADPH oxidase activation in mouse macrophages and magnitude of inflammatory responses in vivo. J Leukoc Biol *101*, 449-457. 10.1189/jlb.3AB0316-139R.
5. Bagaitkar, J., Matute, J.D., Austin, A., Arias, A.A., and Dinauer, M.C. (2012). Activation of neutrophil respiratory burst by fungal particles requires phosphatidylinositol 3-phosphate binding to p40(phox) in humans but not in mice. Blood *120*, 3385-3387. 10.1182/blood-2012-07-445619.
6. Belambri, S.A., Marzaioli, V., Hurtado-Nedelec, M., Pintard, C., Liang, S., Liu, Y., Boussetta, T., Gougerot-Pocidalo, M.-A., Ye, R.D., Dang, P.M.-C., and El-Benna, J. (2022). Impaired p47phox phosphorylation in neutrophils from patients with p67phox-deficient chronic granulomatous disease. Blood *139*, 2512-2522. 10.1182/blood.2021011134.
7. Abe, T., and Hajishengallis, G. (2013). Optimization of the ligature-induced periodontitis model in mice. J Immunol Methods *394*, 49-54. 10.1016/j.jim.2013.05.002.
8. Pandey, S.D., Perpich, J.D., Stocke, K.S., Mansfield, J.M., Kikuchi, Y., Yakoumatos, L., Muszyński, A., Azadi, P., Tettelin, H., Whiteley, M., et al. (2023). Impact of Polymicrobial Infection on Fitness of Streptococcus gordonii <i>In Vivo</i>. mBio *14*, e00658-00623. doi:10.1128/mbio.00658-23.
9. Tseng, J.C., and Kung, A.L. (2012). In vivo imaging of inflammatory phagocytes. Chem Biol *19*, 1199-1209. 10.1016/j.chembiol.2012.08.007.
10. Jin, S., Singhal, R., Luo, J., Higgins, Y.H., Cooper, K.N., Terekhova, M., Carey, K.A., Duff, A.F., Artyomov, M.N., Jala, V.R., et al. (2026). The Nox2 NADPH oxidase regulates neutrophilic inflammation in the oral cavity. Mucosal Immunology *19*, 100341. https://doi.org/10.1016/j.mucimm.2026.04.002.
